# The emotional valence and features of subliminal priming images impact conscious perception of face expressions

**DOI:** 10.1101/255489

**Authors:** Melissa A. Huang, Kunjan D. Rana, Lucia M. Vaina

## Abstract

We investigated, in young healthy subjects, how the affective content of subliminally presented priming images and their specific visual attributes impacted conscious perception of facial expressions. The priming images were broadly categorised as aggressive, pleasant, or neutral and further subcategorised by the presence of a face and by the centricity (egocentric or allocentric vantage-point) of the image content. Subjects responded to the emotion portrayed in a pixelated target-face by indicating via key-press if the expression was angry or neutral. Priming images containing a face compared to those not containing a face significantly impaired performance on neutral or angry target-face evaluation. Recognition of angry target-face expressions was selectively impaired by pleasant prime images which contained a face. For egocentric primes, recognition of neutral target-face expressions was significantly better than of angry expressions. Our results suggest that, first, the affective primacy hypothesis which predicts that affective information can be accessed automatically, preceding conscious cognition, holds true in subliminal priming only when the priming image contains a face. Second, egocentric primes interfere with the perception of angry target-face expressions suggesting that this vantage-point, directly relevant to the viewer, perhaps engages processes involved in action preparation which may weaken the priority of affect processing.

## Introduction

Faces are a rich source of visual information. Within a fraction of a second, humans can recognise emotion, sex, relative age, race, and identity, just by viewing a face. In 1872, Charles Darwin proposed that facial expressions are evolved behaviours that have a biologically adaptive function. Nearly one hundred years later, Paul Ekman (1970) defined six universal facial expressions that are recognised and expressed cross-culturally—happy, sad, fear, disgust, surprise, and anger. Facial expressions are indicators of underlying emotional states, and recognition of the emotion in facial expressions is critical to interpersonal communication, response to imminent threat, and social behaviour. There is substantial evidence that the emotion of facial expressions can even be processed in absence of conscious awareness (see Tamietto and de Gelder (2010) for a review). Functional neuroimaging studies have indicated that non-consciously perceived emotional facial expressions activate subcortical structures (Whalen et al.,1998). De Gelder, Vroomen, Pourois, and Weiskrantz (1999) reported on a blindsight patient who could discriminate among facial expressions presented in his blind field. While most studies indicate that the affect of emotional facial expressions induces neurophysiological changes and influences behaviour without being consciously perceived, there is recent evidence that affective processing does not occur outside of awareness and that any unconscious reaction is dependent on prior semantic processing (Lähteenmäki, Hyönä, & Koivisto, 2015; Nummenmaa, Hyönä, Calvo, 2010).

Major contributions to evidence that affective information can be processed without awareness comes from affective priming experiments. The affective primacy hypothesis (Zajonc, 1980) proposes that affective information can be accessed automatically, and precedes conscious cognition. Subliminal affective priming experiments have provided both neural and behavioural evidence that affective stimuli are unconsciously processed (Li, Zinbarg, Boehm, & Paller, 2008; Whalen et al., 1998). Subliminally presented affective information has also been shown to influence the evaluation of emotionally ambiguous facial expressions (Lee, Kang, Lee, Namkoong & Kyoon An, 2011).

Numerous subliminal priming experiments (Klinger, Burton, & Pitts, 2000; Neumann & Lozo, 2012; Otten & Wentura, 1999), showed that affectively congruent prime-target pairs facilitate either response time or evaluation of the target stimulus. This is referred to as the congruency effect. On the other hand, a study by Hermans, Spruyt, De Houwer, and Eelen (2003) reports the opposite effect on response time (affectively congruent trials were slower than incongruent trials) and Andrews, Lipp, Malan, and Könjg (2011) reported lack of congruent affective priming for subliminal trials, thus providing evidence against the congruency effect.

The discrepancy between these studies may be due to the content and the type of the prime and target stimuli that have been used. The current literature makes little distinction between emotional stimuli that contain faces versus those that do not, although there is convincing evidence that faces are processed separately from non-face stimuli (Kanwisher, McDermott, & Chun, 1997). A large number of affective subliminal priming studies employ facial expressions as stimuli (Aguado, Garcia-Gutierrez, Castañeda, & Saugar, 2007; Andrews, Lipp, Mallan, & Könjg, 2011; Haneda, Nomura, Iidaka, & Ohira, 2003; Wagenbreth, Rieger, Heinze, & Zaehle, 2014) beacuse they are innately association with emotions. However, there are important differences between affective stimuli with and without faces. It has been reported that affective facial expressions are preferentially processed over competing affective non-face visual stimuli perceived either consciously (Batty & Taylor, 2003; Johnson, 2005) or unconsciously (Öhman, 2002). A recent study by Wentura, Rohr, and Degner (2017) shows that, for a short SOA (43 ms), specific negative emotions (angry versus fearful or sad) had distinct impacts on affective priming.

In a preliminary study of schizophrenic subjects (Vaina, Rana, Cotos Cheng, Huang, & Podea, 2014), we investigated the effect of affective subliminal priming on the evaluation of facial expressions and found that subliminal affective priming impacted facial expression recognition in patients with high scores on PANSS positive (indicating delusion, suspiciousness, and hostility) and YMRS (indicating mania) scales, but not the HAM-D (indicating depression) scale. Using the same experimental protocol as in our previous study (Vaina et al., 2014), here we investigate in young healthy subjects how the affective content of the priming image, the presence of a face or the vantage-point of the prime content (image centricity) impact the effect of affective subliminal priming on the identification of target-face expressions.

## Methods

The experimental design replicated that of our previously published paper (Vaina et al., 2014). In the present study, we aimed to investigate the effects of different classes of priming images on the perception of facial expressions in young healthy subjects.

### Subjects

We determined that a sample size of at least 28 subjects would be sufficient to reach a power of 0.80, from an a priori power analysis using G*Power. Thirty-three subjects (17 females, mean age = 18.17 years, SD ± 1.49) participated in the study. All participants were right-handed, had normal or corrected to normal vision, and none had a history of neurologic, psychiatric, or developmental disease. Written consent was given by all participants before the start of the experimental sessions in accordance with Boston University’s Institutional Review Board Committee on research involving human subjects. All data were collected at the Brain and Vision Research Laboratory, Boston University, Boston Massachusetts.

### Experimental procedures

The details of the experimental procedures are presented in Supplement A. Subjects were administered three tests in the following order: SAFFIMAP (Subliminal Affective Image Priming), PV (Prime Validation), and FEP (Face Emotion Processing) tests. The SAFFIMAP test is described below, and the PV and FEP tests are described in Supplement C. For the SAFFIMAP test, both percent correct (described below) and reaction time (shown in Supplement D) were acquired.

### Test Stimuli

The priming images used in this experiment were categorised by their affect (aggressive, pleasant, or neutral), and further sub-categorised by the vantage-point of the image content (egocentric or allocentric) and the presence or absence of a face. The details of the generation and selection of test stimuli are presented in Supplement B.

### Subliminal Affective Image Priming Task (SAFFIMAP)

The SAFFIMAP test assesses the effect of subliminally-presented affective priming images on the perception of a target-face expression (neutral or angry). A schematic view of the experimental trial is presented in Figure 1. First, subjects were asked to press any key on the computer keyboard when prompted by an exclamation mark (6 degrees in height) displayed in the center of the computer screen. This was followed by a 500 ms blank screen at a neutral gray intensity (42 cd/m^2^). Next, a priming image, randomly selected from the experimental database, was shown for 16 ms and immediately followed by a mask displayed for 100 ms. Both the prime and the mask subtended 12.5x12.5 degrees, and were presented in the same spatial location at the center of the computer screen. The target-face was displayed for 500 ms. Subjects responded by pressing a predetermined key on the computer keyboard to report if they perceived the expression of the target-face as neutral or angry. The next trial began immediately after the subject’s response, or after 4000 ms, if no response was entered. There were in total 120 trials. Each contained a unique prime image and a corresponding mask, while the target-faces were randomly selected from 56 face expressions (28 angry and 28 neutral).

**Figure 1.**
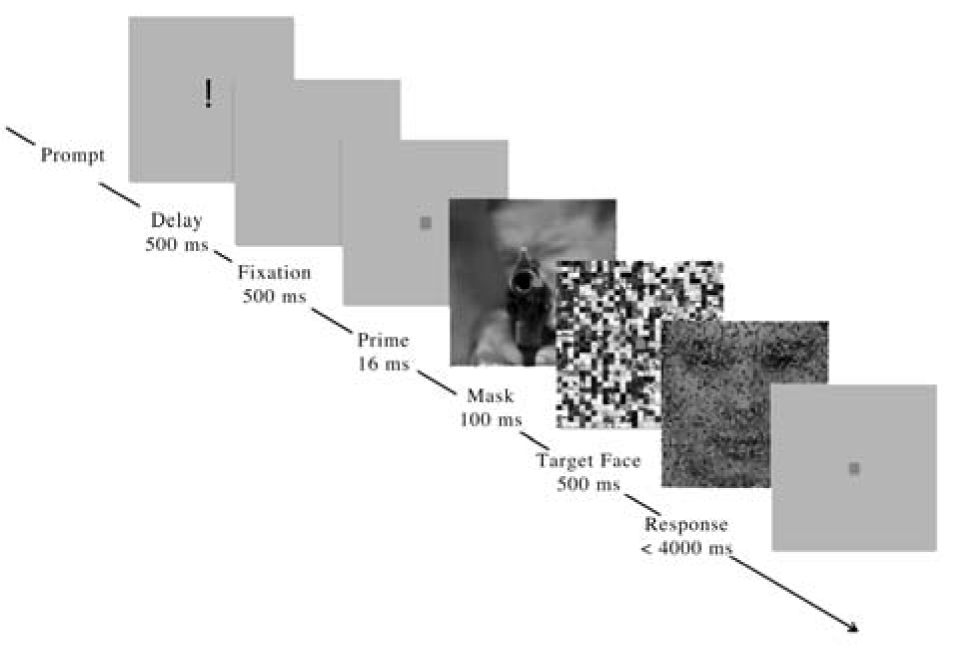
Timeline of a single trial in the SAFFIMAP test.

After completing the test all subjects reported that they were unaware of the priming images.

### Analysis Methods

We applied a General Linear Mixture Model (GLMM) to model the effect of the priming categories (affective category, face content (face present vs. face absent), and centricity) and the target-face (angry vs. neutral) on the evaluation of the target-face expression via a probit-link function, due to the binary response variable (correct vs. incorrect). Subject variability was treated as a random effect in the GLMM. Post hoc tests were conducted through the Wilcoxon Signed-Rank (WSR) Test, where pair-wise comparisons were conducted among average performance within categories between each subject. Significance scores from WSR were FDR-corrected at a false discovery rate of 0.05. Although all priming category and target face interactions were analyzed, only those proved significant by the GLMM analysis are reported below.

## Results

### 1. Overall target face expression recognition with face and non-face primes

The presence or absence of a face in the priming images significantly affected subjects’ evaluation of the target-face (GLMM, F(1,2976) = 6.186, p = 0.013)). In trials where primes contained a face, subjects were poorer at correctly identifying the emotion expressed in the target face, compared to when the prime image did not contain a face. (face: μ = 74.20%, se = 1.79%; non-face: μ = 78.00%, se = 1.49%) (WRS, z=−2.266, p = 0.023)

This effect differed significantly depending on the affective valence of the priming image (F(2,2976) = 5.805, p = 0.003). Only primes containing affective facial expressions (aggressive and pleasant) contributed to the detrimental effect of face primes compared to non-face primes on the evaluation of target-face expression. The difference in performance observed when the prime contained a face versus when it did not was significant only when the affective content of the prime image was aggressive (aggressive, face: μ = 73.4%, se = 1.99%; aggressive, non-face: μ = 81.0%, se = 1.85%) (WRS, z = −3.835, p = 0.001). When the prime image was pleasant, the effect of the face content category approached significance (pleasant, face: μ = 72.4%, se = 2.68; pleasant, non-face: μ = 76.4%, se = 2.26) (WRS, z = −1.559, p = 0.132) but when the prime image was neutral, the effect was not significant (neutral, face: μ = 76.8%, se = 1.58%; neutral, non-face: μ = 76.6%, se = 1.97%) (WRS, z = 0.104, p =0.918).

#### 1.2. The effect of the emotional content in a face prime on target-face expression recognition

The difference in target-face expression evaluation between when priming with neutral, face-containing images versus aggressive or pleasant face-containing images was significant. Subjects performed better on perceiving target-face expressions when the affective value of the face-containing prime image was neutral compared to aggressive or pleasant (neutral vs. aggressive: WRS, z = 2.060, p = 0.039) (neutral vs. pleasant: WRS, z = 1.966, p = 0.049). There was no significant statistical difference between priming with pleasant or aggressive face images (WRS, z = −0.248, p = 0.804). There was no significant statistical difference between the affective categories for primes which did not contain a face.

### 2. Neutral expression recognition with face and non-face primes

For evaluating specifically neutral expressions when primed with aggressive images, performance was significantly better when the prime did not contain a face compared to when it contained a face (WRS, z = −3.054, p = 0.0022) (aggressive, face: μ = 74.53%, se = 0.028%). In addition, performance was significantly better when the non-face prime image was aggressive (μ = 85.33%, se = 2.11%) compared when it was neutral (μ = 78.97%, se = 2.46%) (WRS, z = −2.426, p = 0.015)

### 3. Angry expression recognition with face and non-face primes

Performance on perceiving angry target-face expressions when primed with a face-containing image was significantly lower when the prime was pleasant (μ = 64.21%, se = 4.18%) compared to when the prime was neutral (μ = 73.60%, se = 3.58%) (WRS, z = 2.449, p = 0.014).

### 4. The effect of centricity of the primes on target-face expression recognition

We found a significant effect of the vantage-point of the priming image and the target-face expression (GLMM, F(1,2976) = 0.041). When evaluating neutral target-faces, subjects were more often correct when the vantage-point of the priming image was egocentric compared to when it was allocentric (egocentric: μ = 81.64%, se = 2.03%; allocentric: μ = 77.31%, se = 2.05%) (WRS, z = -2.215, p = 0.017). The reverse effect on performance was seen when evaluating angry target-faces, although the difference was not statistically significant. There was slightly lower performance for egocentric priming image centricity (μ = 69.62%, se = 2.82%) than for allocentric centricity (μ = 72.26%, se = 3.22%). Subjects also more often evaluated the target-faces as neutral compared to angry when the priming image was egocentric (egocentric prime, angry target-face: μ = 69.62%, se = 2.82%; egocentric prime, neutral target-face: μ = 81.64%, se = 2.03%) (WRS, z = -2.935, p = 0.003).

**Figure 2.**
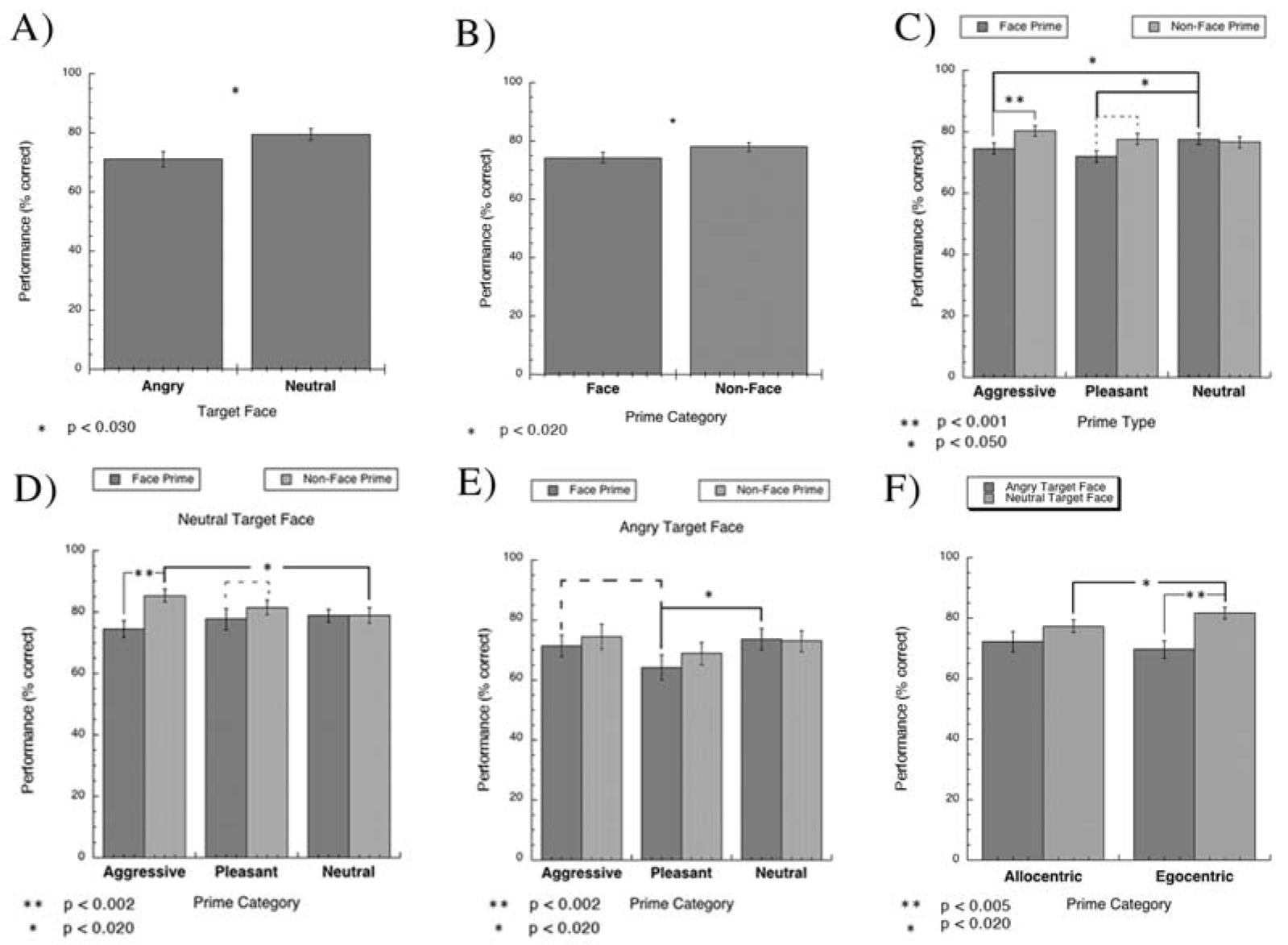
Performance Results. A) Performance was significantly higher for identifying neutral facial expressions compared to angry facial expressions. *B)* Overall performance was significantly worse when the priming image contained a face compared to when it did not contain a face. *C)* Performance was only impaired by primes containing faces when the priming image was also affective. In addition, when the priming image contained a face, performance was worse when the prime was pleasant or aggressive compared to when the prime was neutral. *D)* For evaluating specifically neutral expressions when primed with aggressive images, performance was significantly better when the prime did not contain a face compared to when it contained a face. In addition, performance was significantly better when the non-face prime image was aggressive compared when it was neutral. *E)* Performance on perceiving angry target-face expressions when primed with a face-containing image was significantly lower when the prime was pleasant compared to when the prime was neutral. *F)* When evaluating neutral target-faces, subjects were more often correct when the vantage point of the priming image was egocentric compared to allocentric. Subjects also more often evaluated the target-face as neutral compared to angry when the priming image was egocentric

## General Discussion

Our first notable finding is that subliminally priming with a face-containing image significantly impairs the subjects’ ability to perceive the emotional expression of the target-face. This suggest that processing of faces occurs preferentially to the processing of non-face containing images, and that the face perceived in the priming image interferes with the ability to perceive the target-face expression. This result is in agreement with Kemps, Erauw, and Vandierendock (1996) finding that facial expressions influenced the liking rating of Korean ideographs at exposure duration of 30 ms and longer, while pictures without faces did not elicit an affective reaction until they were presented for at least 100 ms. Consistent with their results, we suggest that the affective information in an image containing a face is processed more immediately than the affective information in an image not containing a face.

Furthermore, we showed that a prime containing a pleasant or aggressive face expression elicits significantly poorer performance compared to a prime with a neutral face expression, regardless of the target-face expression. This result is at odds with the findings of Andrews et al. (2011), who reported that there was no subliminal affective priming effect when priming with emotional facial expressions (Andrews, Lipp, Malan, & Könjg, 2011). Our result suggests that the emotional expression of a face in a prime can be processed, at least partially, during a 16 ms exposure, and that this interferes with the conscious processing of the target-face expression. Taken together these results suggest that the face in the prime image prompts the viewer to expect an emotionally charged target-face, which quickens (see Supplement D) and biases the response towards the angry rather than neutral-target face. Consistent with studies reporting an affective congruency effect with subliminal primes (Klinger, Burton, & Pitts, 2000; Neumann & Lozo, 2012; Otten & Wentura, 1999), we found that specifically the incongruency of the pleasant prime and the angry expression of the target-face lead to worse performance compared to priming images containing neutral or aggressive face expressions. Differing from these studies, we found that this effect occurred only when the prime image contained a face.

Regardless of whether the prime image contained a face, there was a significant effect of the centricity of the image on the evaluation of the target-face expression. Subjects were more likely to evaluate the facial expression as neutral when the content of the prime was egocentric. Furthermore, priming neutral target-faces with egocentric images resulted in significantly better performance compared to priming with allocentric images. These results show that the vantage-point of the subliminally presented prime image (16 ms presentation) affects the perception of emotional expression of the target-face. From this result, we suggest that, because of the direct relevance of the egocentric vantage-point to the viewer, the egocentric prime image may engage a series of processes relevant to potential action preparation, which could weaken the priority of affective processing, and thus interfere with the ability to recognize the angry target-face expression. Our results suggest that the content of the priming image (presence of a face) and its vantage-point (egocentric or allocentric) impacts the effect of affective subliminal priming on the evaluation of facial expressions. In this study, our interest was to investigate the effect of multiple priming categories on two distinct target-face expressions (angry and neutral). We will extend this study by adding additional dimensions to the priming stimuli, to dissociate between the emotion of the scene and that of the facial expressions. Furthermore, we will use fMRI to investigate the neuronal substrate of different dimensions of subliminal affective priming in healthy subjects and clinical populations.

## Acknowledgments

The authors thank all subjects for participating in this research. We thank Leon Chen for helping with stimuli generation.

